# Prokaryotic nanocompartments form synthetic organelles in a eukaryote

**DOI:** 10.1101/244095

**Authors:** Yu Heng Lau, Tobias W. Giessen, Wiggert J. Altenburg, Pamela A. Silver

**Author notes:** Y.H.L. and T.W.G. contributed equally. Y.H.L., T.W.G. and P.A.S. designed the research and wrote the manuscript. Y.H.L., T.W.G. and W.J.A. performed the experiments. **Corresponding author:** Prof. Pamela Silver.

## Abstract

Compartmentalization of proteins into organelles is a promising strategy for enhancing the productivity of engineered eukaryotic organisms. However, approaches that co-opt endogenous organelles may be limited by the potential for unwanted crosstalk and disruption of native metabolic functions. Here, we present the construction of synthetic non-endogenous organelles in the eukaryotic yeast *Saccharomyces cerevisiae*, based on the prokaryotic family of self-assembling proteins known as encapsulins. We establish that encapsulins self-assemble to form nanoscale compartments in yeast, and that heterologous proteins can be selectively targeted for compartmentalization. Housing destabilized proteins within encapsulin compartments affords protection against proteolytic degradation *in vivo*, while the interaction between split protein components is enhanced upon co-localization within the compartment interior. Furthermore, encapsulin compartments can support enzymatic catalysis, with substrate turnover observed for an encapsulated yeast enzyme. Encapsulin compartments therefore represent a modular platform, orthogonal to existing organelles, for programming synthetic compartmentalization in eukaryotes.

## Introduction

Intracellular compartmentalization is a fundamental strategy used by all organisms to organize and optimize their metabolism. Examples of compartments in nature range from eukaryotic lipid-bound organelles to prokaryotic protein-based containers^1–4^, and their functions include sequestering toxic metabolic products, generating distinct biochemical environments, and stabilizing otherwise unstable proteins and biosynthetic intermediates. The ability to incorporate similar functional properties in engineered organisms could lead to significant improvements in metabolic engineering and recombinant protein expression^5,6^. However, efforts to reprogram naturally-occurring compartments for synthetic applications are challenging due to their inherent complexity and the large number of different biomacromolecules involved^7–10^. We therefore identified the encapsulin family of self-assembling prokaryotic proteins as a highly engineerable candidate suitable for designing programmable synthetic organelles in eukaryotes^11–14^.

Encapsulins are 25–40 nm diameter hollow compartments comprised of 60 or 180 copies of a single self-assembling capsid protein^11,12^. The varied native functions of encapsulins all involve packaging proteins within their interior as part of the self-assembly process to tailor the activity of packaged components. *In vivo* protein encapsulation is guided by short targeting peptides (TPs), which are located at the *C*-termini of cargo proteins. A large variety of native cargo proteins has been identified in bacteria and archaea, including peroxidases and ferritin-like proteins involved in stress response pathways^11,14–16^. Using *Escherichia coli* as a host, it has been shown that packaging of non-native proteins into the encapsulins from *Thermotoga maritima* and *Brevibacterium linens* can be achieved by fusion of targeting peptides to the intended cargo^17,18^.

Given their modularity and programmability, encapsulins are an ideal platform for building synthetic compartmentalization in eukaryotes. In contrast to approaches that leverage existing organelles^19–22^, encapsulins have the advantage of being completely orthogonal to endogenous eukaryotic compartments. There is also ample choice of different encapsulin protein variants derived from different families of bacteria and archaea. In particular, the encapsulin from *Myxococcus xanthus* has been structurally characterized, and has the ability to simultaneously package three different proteins in its native form^14^.

Here, we present the construction of synthetic organelles in the yeast *Saccharomyces cerevisiae*, based on the *Myxococcus xanthus* encapsulin^14^. We show that encapsulin compartments can stabilize heterologous cargo proteins against degradation, co-localize proteins within their interior, and act as nanoreactors for housing enzymatic catalysts (Fig. 1A). In doing so, we demonstrate that protein-based compartments can mimic the ability of eukaryotic organelles to control protein localization and activity in living cells.

**Figure 1.**
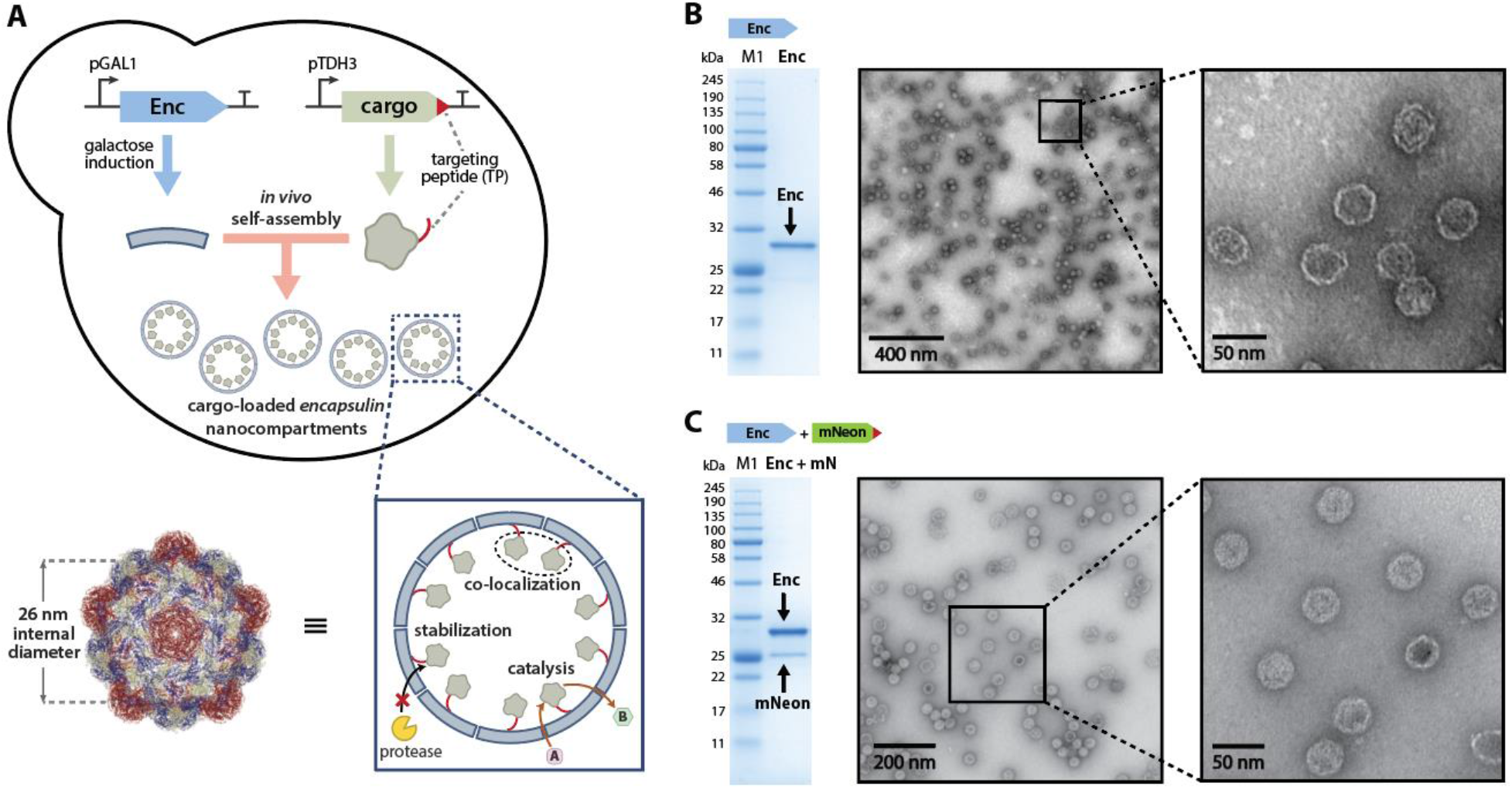
A) Co-expression of encapsulin and targeted cargo in yeast results in self-assembly of cargo-loaded nanocompartments, with an internal diameter of 26 nm (structure from PDB: 4PT2). Encapsulin compartments can stabilize and co-localize cargo proteins, as well as allow for catalysis to occur within their interior. B) Encapsulin EncA (32.5 kDa) can be purified to homogeneity from yeast, as determined by SDS-PAGE and TEM. C) Heterologous proteins such as mNeonGreen (28.4 kDa) can be packaged inside encapsulin compartments, as determined by SDS-PAGE and TEM on the co-purified sample.

## Results

### Self-assembly of cargo-loaded encapsulin compartments in yeast

Expression and self-assembly of encapsulin compartments in yeast was achieved using a plasmid containing the encapsulin gene EncA from *M. xanthus*^14^ under control of the inducible GAL1 promoter (Fig. 1B). A clear induction band was observed by SDS-PAGE for cultures grown in galactose induction media, corresponding to the 32.5 kDa encapsulin monomer (SI Fig. S3.3). The identity of the band was confirmed to be the EncA by mass spectrometry. Isolation of the encapsulin compartments was achieved by PEG precipitation from the cell lysate, followed by size-exclusion and ion-exchange chromatography, resulting in a pure sample of encapsulin as determined by SDS-PAGE (Fig. 1B and SI Fig. S3.4). Under native PAGE conditions, the size of the encapsulin particle was >1 MDa, consistent with the formation of a self-assembled capsid structure (SI Fig. S3.5). Transmission electron microscopy (TEM) of negatively-stained samples confirmed that the purified encapsulins were highly homogeneous spherical capsids with the expected diameter of 32 nm^14^ (Fig. 1B and SI Fig. S4.1).

*In vivo* self-assembly of protein cargo inside encapsulins was demonstrated using the fluorescent protein mNeonGreen with a C-terminal fused targeting peptide (TP, sequence PEKRLTVGSLRR) under control of the constitutive TDH3 promoter (Fig. 1C). After encapsulin induction and subsequent purification of the capsids from yeast, co-purification of mNeonGreen with encapsulin was observed by SDS-PAGE (Fig. 1C). The cargo loading percentage relative to encapsulin was estimated to be 24% (~43 molecules per compartment) based on gel densitometry. Fluorescence was also confirmed to be associated with the assembled capsids by in-gel fluorescence of the purified encapsulin band on a native PAGE gel (SI Fig. S3.6). Confirmation that the cargo-loaded encapsulins had assembled as expected was obtained by TEM (Fig. 1C). Furthermore, the purified encapsulins were remarkably stable over time, with minimal deterioration observed by native PAGE and TEM despite storage for two months at 4 °C in Tris buffer (SI Fig. S5.1).

### Protection of encapsulated cargo from degradation *in vivo*

Cargo proteins packaged inside encapsulin compartments were protected against proteolytic degradation (Fig. 2). A destabilized cargo protein was created by appending mNeonGreen with a *C*-terminal CLN2-PEST degradation tag, followed by the targeting peptide for encapsulation. Yeast cells expressing this unstable fusion protein (mN-PEST-TP) only showed high levels of *in vivo* fluorescence when co-expressed with encapsulin, as determined in bulk measurements (Fig. 2B) and by fluorescence microscopy of individual cells (Fig. 2C). Minimal fluorescence intensity was observed when the TP was removed or the encapsulin was not induced. Based on bulk plate-reader fluorescence measurements (Fig 2D), an 11-fold increase was observed for cargo protein levels as a result of encapsulation.

**Figure 2.**
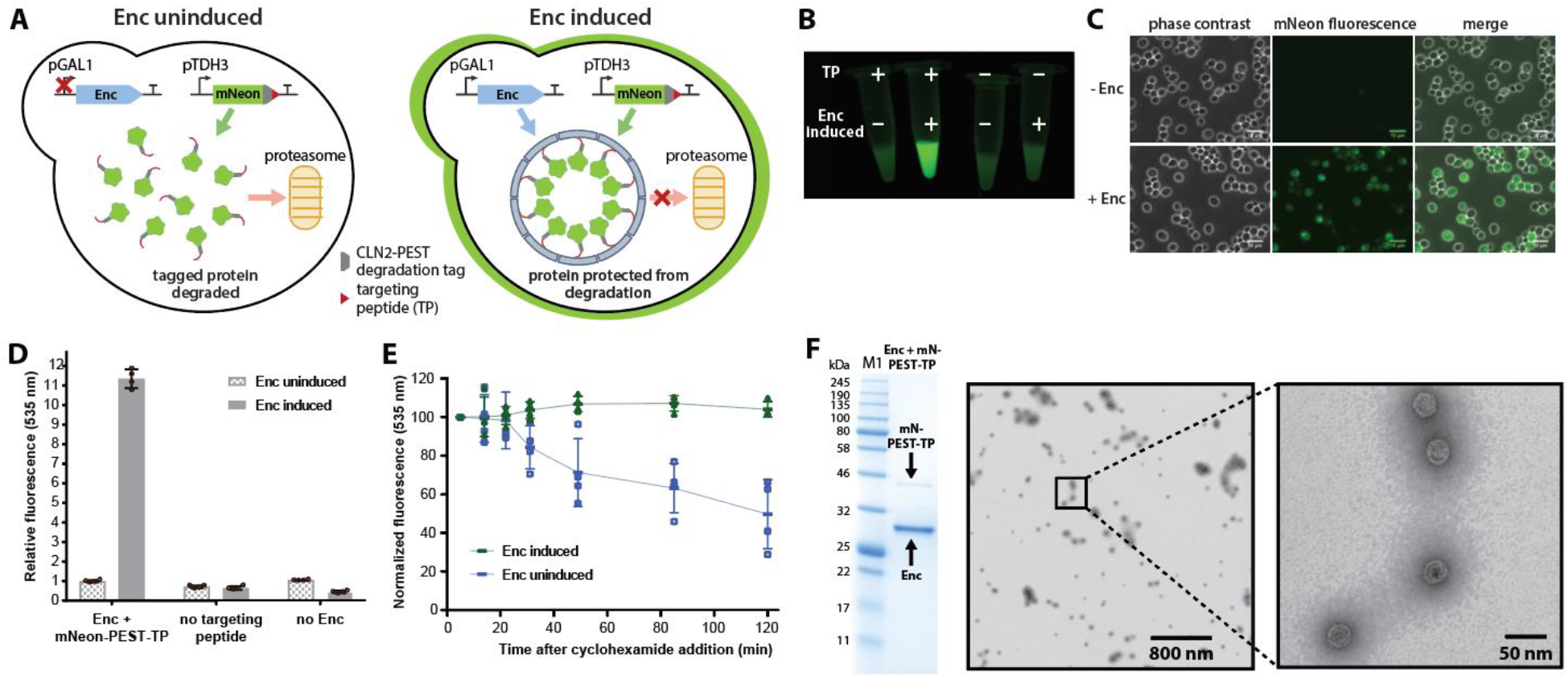
A) Encapsulation of cargo proteins can enhance stability against proteolytic degradation. Destabilized mNeonGreen bearing a PEST degradation tag shows an 11-fold increase in *in vivo* lifetime as determined by B) the fluorescence intensity in bulk images, C) imaging of cells by fluorescence microscopy, and D) bulk plate reader fluorescence intensity measurements. E) Upon inhibiting protein synthesis using cyclohexamide, destabilized mNeonGreen was protected from degradation only when encapsulated. F) Isolation of the loaded encapsulins from yeast showed co-purification with destabilized mNeonGreen (47.6 kDa) by SDS-PAGE and TEM.

The stabilization effect of encapsulation was also observed after inhibition of new protein synthesis (Fig 2E). Upon inhibition using cyclohexamide, yeast cells expressing only the destabilized cargo showed a gradual loss of fluorescence over 2 h. In comparison, yeast cells co-expressing the cargo and encapsulin maintained a constant level of fluorescence over the 2 h period. To confirm the integrity of the encapsulin compartments, the loaded compartments were co-purified from yeast as before, displaying associated fluorescence by PAGE, and proper assembly by TEM (Fig. 2F and SI Fig. S3.7).

### Co-localization of split protein components within encapsulins

Multiple heterologous proteins can be co-localized inside encapsulin compartments (Fig. 3A). Using an established split-Venus system^23^, an elevated level of fluorescence was only observed when both split components were targeted for encapsulation, and the encapsulin gene itself was present and induced (Fig. 3B-D). A 2.5-fold increase in fluorescence intensity was observed, consistent with the fluorescence response previously reported when the split components are co-localized^23^ (Fig. 3C). Co-encapsulation of two distinct proteins did not disturb encapsulin assembly as indicated by the high molecular weight band on native PAGE (Fig. 3E) and the readily assembled particles observed using TEM (Fig. 3F). The split components were estimated to have a cargo loading percentage of 42% for Ven-N and 30% for Ven-C (~76 and 54 per compartment respectively) based on SDS-PAGE analysis (Fig. 3F).

**Figure 3.**
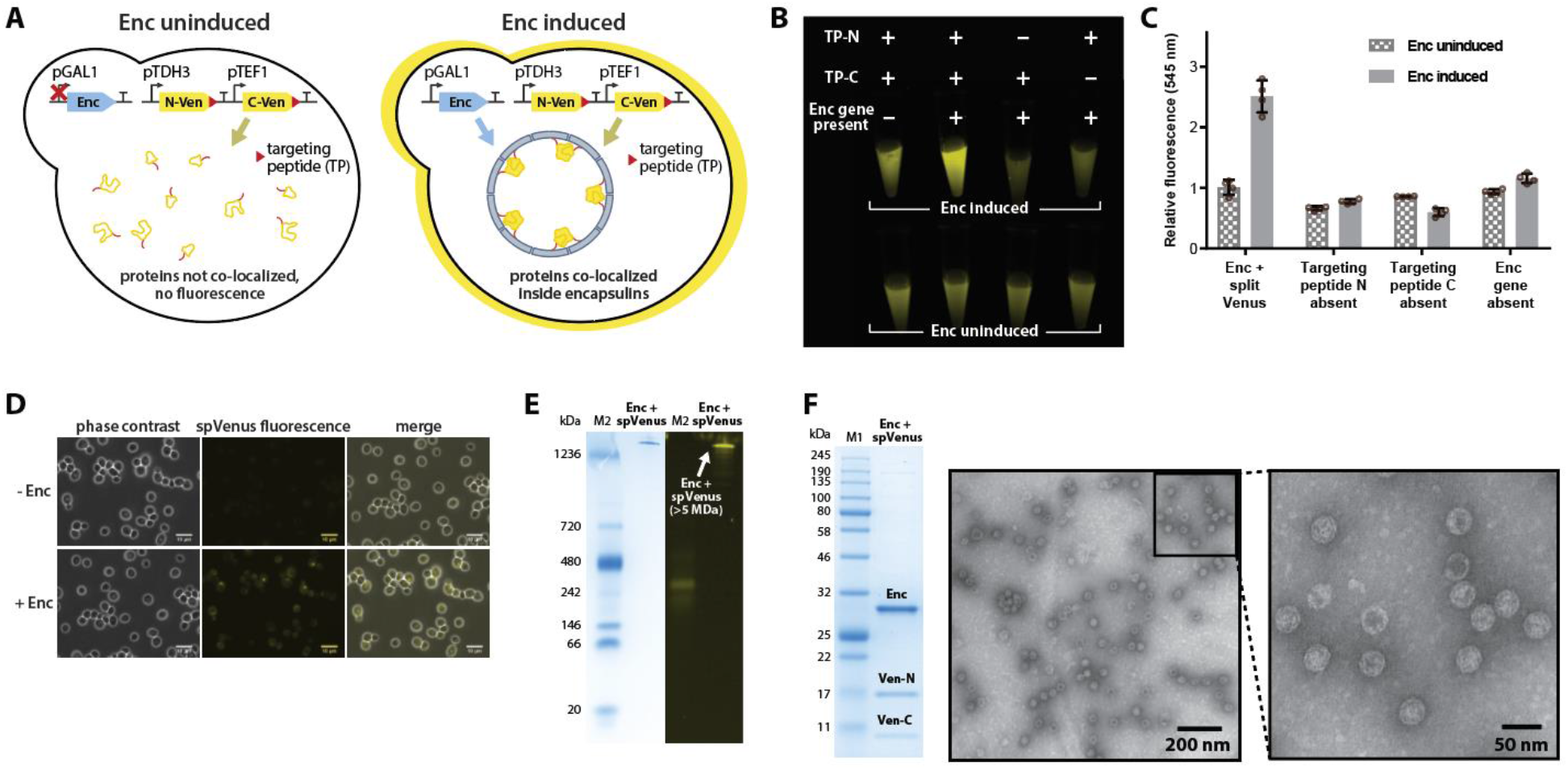
A) Co-encapsulation of split-Venus components led to an increase in fluorescence intensity as determined by B) bulk images, C) bulk plate-reader measurements, and D) fluorescence microscopy. E) Native PAGE analysis of encapsulated split-Venus showed fluorescence associated with the high molecular weight assembled encapsulin nanocompartment band. F) Purified encapsulins co-purified with the split-Venus proteins (19.4/11.6 kDa) by SDS-PAGE, and formed the expected compartments as imaged by TEM.

### Catalytic turnover of an encapsulated yeast enzyme

Finally, enzyme-loaded encapsulins were shown to be viable nanoreactors for catalytic processes. The assembled encapsulin structure^14^ contains small 5–10 Å sized pores, which in principle, can allow small molecule substrates and products to diffuse in and out of the compartment. The candidate enzyme chosen for encapsulation was Aro10p, a tetrameric pyruvate decarboxylase enzyme that is endogenous to yeast, participating in the catabolism of aromatic amino acids such as tyrosine (Fig. 4A). In particular, Aro10p catalyzes the decarboxylation of 4-hydroxyphenylpyruvate (4-HPP) to 4-hydroxyphenylacetaldehyde (4-HPAA)^24^. There is great interest in the production of 4-HPAA in yeast, as its reaction with dopamine *via* Pictet-Spengler cyclization leads to norcoclaurine, a key intermediate for the heterologous production of many valuable medicinal benzylisoquinoline alkaloids of the opioid family^25–27^. Two challenges associated with 4-HPAA production in yeast are instability due to endogenous aldehyde and alcohol dehydrogenases, and toxicity effects associated with aldehyde reactivity^28,29^. We sought to test if Aro10p could be encapsulated within encapsulins as a potential route towards addressing these challenges.

**Figure 4.**
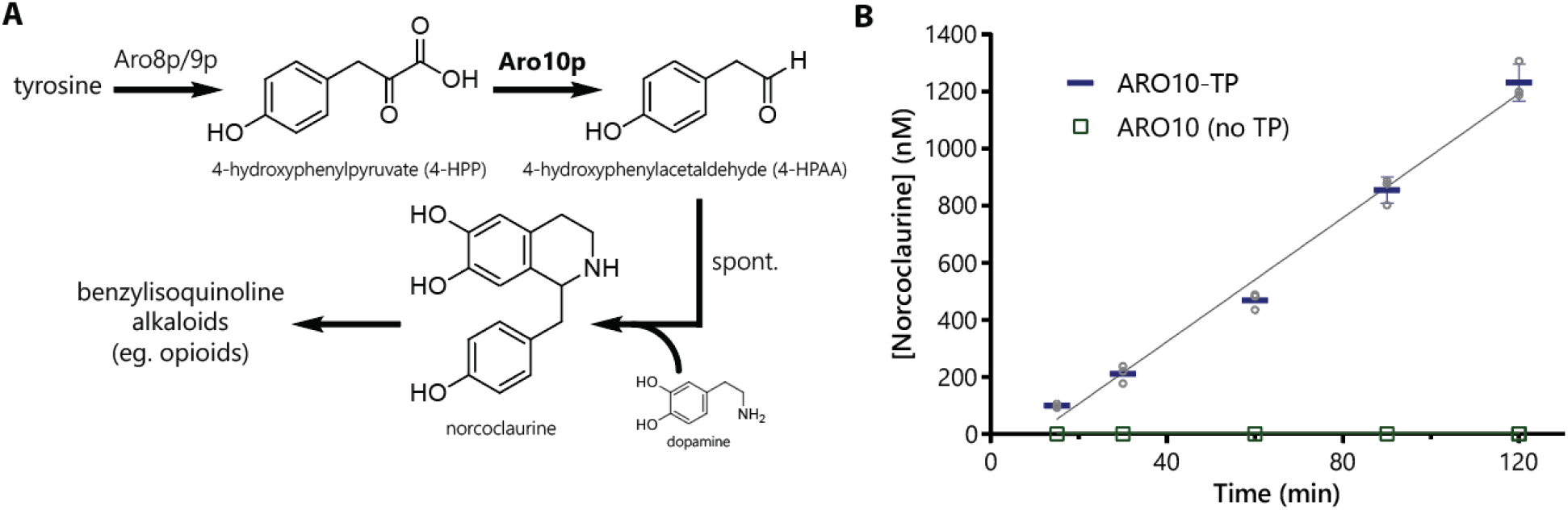
A) The Aro10p enzyme is involved in tyrosine catabolism, generating 4-hydroxyphenylacetaldehyde (4-HPAA) which can be measured by QTOF-LCMS after conversion to norcoclaurine by spontaneous reaction with dopamine. B) Encapsulin nanocompartments show enzymatic activity only when co-expressed with the version of Aro10p that is fused with the targeting peptide (TP).

Encapsulin nanoreactors containing the Aro10-TP enzyme displayed enzymatic 4-HPP decarboxylation activity (Fig. 4B). Comparing the encapsulins purified from yeast strains co-expressing either Aro10-TP or Aro10 with no TP, only the encapsulins co-expressed with Aro10-TP showed *in vitro* enzymatic activity in the presence of 4-HPP. Enzymatic activity of purified encapsulins was determined by spontaneous Pictet-Spengler cyclization of the 4-HPAA product with dopamine to give norcoclaurine, which could then be detected by QTOF-LCMS (SI Fig. S7.1). To confirm the fidelity of the targeting process, the purified encapsulins with TP or no TP were compared by SDS-PAGE, indicating the presence of Aro10p only when the TP was present (SI Fig. S3.8).

## Discussion

In this work, we establish encapsulins as a platform for engineering synthetic compartmentalization in yeast. There are several key features of encapsulins that distinguish them from other related compartmentalization systems currently being studied. The first is the ability to self-assemble with its associated cargo *in vivo*, using only a single repeating protein unit and short peptide tag. Other proteinaceous organelles, such as bacterial microcompartments, are comprised of many different protein subunits and thus entail a higher degree of complexity. Although significant progress has been made towards understanding the molecular principles governing these complex systems^1,10^, our incomplete understanding is still a bottleneck for repurposing such systems as synthetic organelles.

The orthogonality of encapsulins in context of eukaryotic organisms is another distinct aspect of our approach to synthetic compartmentalization. Recent reports have explored the localization of engineered proteins to eukaryotic compartments such as the peroxisome^21,22^, mitochondria^19^, and vacuoles^30^. While large scale reprogramming may be tolerated for some organelles, essential functions may be perturbed. The other restriction imposed when using native organelles is that the protein import mechanism, biochemical environment, and substrate permeability may also be difficult to modify. There are examples where the organelle environment is advantageous, such as using the oxidative environment of the mitochondria^19^. In a completely orthogonal synthetic compartment, these parameters can be tailored with much greater freedom.

Encapsulins present a tunable platform for maximizing the productivity of engineered pathways. The encapsulin targeting system enables co-localization of multiple enzymes. Co-localization could lead to significant rate enhancements for reactions involving unstable or toxic intermediates, or in situations where a high local concentration of the intermediate is required^31^. The levels of each enzyme within encapsulins could be controlled by modifying the targeting peptide sequence and hence the strength of its interaction with the encapsulin protein. Substrate accessibility may be tuned by engineering the residues adjacent to the compartment pores. Furthermore, thousands of encapsulin variants exist thereby providing compartments with different diameters, surface charges, pore sizes, and other biophysical properties that can be chosen.

In conclusion, we have shown that encapsulin compartments display many of the properties required for building synthetic organelles in eukaryotes. Encapsulin compartments can extend the lifetime of unstable proteins, and co-localize proteins to induce proximity effects. The encapsulin platform is also capable of serving as a nanoreactor, with encapsulated enzymes displaying catalytic activity. Taken together, the encapsulin system is modular and robust, with potential applications for enhancing protein production and metabolic engineering in yeast. This work now paves the way for future studies on controlling new enzymatic chemistry within encapsulins, and the integration of encapsulin organelles into engineered yeast metabolism.

## Methods

DNA sequences, additional PAGE gels and TEM images, information about Gibson cloning constructs and a list of strains are provided in the Supplementary Information.

### Cloning

All inserts were synthesized as codon-optimized gBlocks (IDT). All plasmids were cloned by Gibson assembly using NEBuilder HiFi DNA Assembly Master Mix (NEB). Plasmids were first cloned in 5-alpha competent *E. coli* (NEB), isolated by miniprep, and then transformed into the CEN.PK2-1D strain of *S. cerevisiae* (Euroscarf) using the high efficiency LiAc/SS carrier DNA/PEG method described by Gietz and Schiestl^32^. Linear constructs were obtained from synthetic gBlocks and transformed directly into CEN.PK2-1D strain of *S. cerevisiae* (Euroscarf) using the high efficiency LiAc/SS carrier DNA/PEG method described by Gietz and Schiestl^32^. Cassettes were directed to disrupt the HO locus of the yeast genome. Selection for the KanMX resistance marker was carried out on G418 plates. Integrants were confirmed by PCR and sequencing of the integrated cassette, starting from regions outside the cassette.

### Encapsulin expression and purification

Overnight 5 mL liquid cultures of yeast strains in synthetic defined dropout media were diluted into 50 mL of fresh media and grown at 30 °C for 18–24 h. Cells were resuspended in 6 mL PBS buffer and lysed using glass beads, and then sodium chloride and PEG-8000 were added to the soluble fraction to a final concentration of 0.5 M and 8% respectively. After sitting for 15 min on ice, the precipitate was isolated, redissolved in 2 mL PBS buffer, and purified by size exclusion using a HiPrep 16/60 Sephacryl S-500 HR column (GE Healthcare) in PBS buffer (1 mL/min) on an AKTA Explorer (Amersham Biosciences). The encapsulin fractions were concentrated using Amicon Ultra-15 Centrifugal Filter Units with Ultracel-100 membrane (Millipore), then diluted in 2 mL of 20 mM Tris buffer at pH 8. Ion-exchange chromatography using a HiPrep DEAE FF 16/10 column (GE Healthcare) resulted in the fully-purified encapsulin sample for further analysis. The gradient used for ion-exchange was as follows: 100% A for 0–100 mL, 100% A to 50% A + 50% B for 100–200 mL, 100% B for 200–300 mL, 100% A for 300–400 mL; where A is 20 mM Tris pH 8, B is 20 mM Tris pH 8 with 1 M NaCl (flow rate: 3 mL/min).

### Polyacrylamide gel electrophoresis

SDS-PAGE was run using Novex WedgeWell 14% Tris-Glycine Mini Gels (Invitrogen), staining with Coomassie Brilliant Blue. Native PAGE was run using NativePAGE™ 3–12% Bis-Tris Protein Gels (Invitrogen), running either under regular Bis-Tris buffer conditions or using NativePAGE running buffers (Invitrogen) for Blue Native PAGE. Color Prestained Protein Standard, Broad Range 11–245 kDa (NEB) was used as a ladder for SDS-PAGE (marked as “M1”), while NativeMark Unstained Protein Standard (Life Technologies) was used for native PAGE (marked as “M2”). Gel images were captured on a ChemiDoc MP Imaging System (Bio-Rad), using the accompanying Image Lab software to approximate band intensities for densitometry measurements. Gel densitometry was carried out using the in-built quantification tools on ImageLab software (Bio-Rad).

### Transmission electron microscopy

Electron microscopy was conducted on a Tecnai G2 Spirit BioTWIN. Encapsulin samples were diluted to approximately 0.1 mg/mL for adsorption onto Formvar carbon coated gold grids 200 mesh FCF200-Au (EMS) after glow discharge. Excess sample was removed by blotting on filter paper (Whatman). Uranyl formate or uranyl acetate was then applied for negative staining.

### Fluorescence measurements

Bulk yeast images - Cells were inoculated into 5 mL of SD-His and grown overnight. After normalizing for OD, the cells were resuspended in 2 mL of water, and 200 μL of this suspension was added to 2 mL of either SD-His or induction media. After 24 h growth at 30 °C, cells were pelleted and resuspended in water (normalizing for OD). Images were taken on a ChemiDoc MP Imaging System (Bio-Rad).

Plate reader measurements - Samples were prepared in the same manner as for bulk imaging, with fluorescence intensity measurements carried out on a Synergy Neo plate reader (BioTek), with excitation/emission wavelengths of 500/535 nm for mNeonGreen and 515/545 nm for Venus. Measurements were carried out in independent biological quadruplicate experiments, each consisting of technical triplicates.

Fluorescence microscopy - Cells prepared for bulk imaging were also imaged directly on a Nikon TE 2000 microscope in glass bottom dishes (MatTek, 35 mm, uncoated, no. 1.5) under agar pad. Microscope light source power, detector gain and image processing settings were kept consistent between images and samples to ensure the validity of any comparative conclusions drawn.

Cyclohexamide chase experiment - Cells were first grown using the same protocol as for bulk imaging described above. From the 24 h induced and non-induced cultures, 1 mL of each culture was pelleted and resuspended into 10 mL of fresh media (OD ~0.8) and grown at 30 °C for 40 min. After this time, 100 μg/uL cyclohexamide was added to each culture, and at each time point, 400 μL aliquots were taken, resuspended in 100 μL H_2_O and snap frozen for later measurement. Fluorescence intensity measurements were obtained by the plate reader method described above. Measurements were carried out in independent biological quadruplicate experiments, each consisting of technical triplicates.

### Enzymatic assays

Enzyme assays were conducted in biological triplicate. The assay conditions were 100 mM potassium phosphate buffer pH 7, 1 mM MgCl_2_, 0.5 mM thiamine pyrophosphate, 1 mM dopamine hydrochloride, 1 mM 4-hydroxyphenylpyruvate and 75 μg/mL encapsulin. Reactions were conducted at 30 °C in 1 mL volumes. At each time point, 100 μL was removed from the reaction and 100 μL of MeCN was added. Any precipitated debris was pelleted, and then 5 μL of the supernatant was added to 45 μL of water to give the final sample ready for QTOF-LCMS analysis.

Norcoclaurine production was measured on a QTOF-LCMS (Agilent 6530), running samples on a Orpak CDBS 453 column (Shodex). The method used for analysis was: 0–9 min 0% B, 9–11 min 0 to 95% B, 11–14 min 95% B, 14–16 min 95 to 0% B, 16–23 min 0% B (flow rate 0.5 mL/min; A = 95% H_2_O + 5% MeCN with 0.1% formic acid, B = 100% MeCN). The MS acquisition parameters were as follows: positive ion mode, gas temperature: 325 °C, drying gas: 10 L/min, nebulizer: 12 psig, VCap: 3500 V, mass range: 100–1000 m/z, acquisition rate: 2 spectra/s, acquisition time: 500 ms/spectrum. Norcoclaurine standards for generating a standard curve were obtained from Toronto Research Chemicals.

## Acknowledgements

Y.H.L. acknowledges funding from the Wellcome Trust (107402/Z/15/Z). T.W.G. was supported by a Leopoldina Research Fellowship (LPDS 2014–05) from the German National Academy of Sciences. This work was supported by the National Science Foundation (NSF) Synthetic Biology of Yeast grant (MCB-1330914).

